# Megasatellite formation and evolution in vertebrates

**DOI:** 10.1101/2021.07.19.452908

**Authors:** Stéphane Descorps-Declère, Guy-Franck Richard

**Affiliations:** Institut Pasteur, Département de Biologie Computationelle, 25 rue du Dr Roux, F-75015 Paris, France; Institut Pasteur, Département Génomes & Génétique, 25 rue du Dr Roux, F-75015 Paris, France; CNRS, UMR3525, F-75015 Paris, France

**Keywords:** Tandem repeat, minisatellite, microsatellite, genomics

## Abstract

Since the formation of the first proto-eukaryotes, more than 1.5 billion years ago, eukaryotic gene repertoire as well as genome complexity has significantly increased. Among genetic elements that are responsible for this increase in genome coding capacity and plasticity are tandem repeats such as microsatellites, minisatellites and their bigger brothers, megasatellites. Although microsatellites have been thoroughly studied in many organisms for the last 20 years, little is known about the distribution and evolution of mini- and megasatellites. Here, we describe the first genome-wide analysis of megasatellites in 58 vertebrate genomes, belonging to 12 monophyletic groups. We show that two bursts of megasatellite formation occurred, one after the radiation between agnatha et gnathostomata fishes and the second one later, in therian mammals. Megasatellites are frequently encoded in genes involved in transcription regulation (zinc-finger proteins) and intracellular trafficking, but also in cell membrane metabolism, reminiscent of what was observed in fungi genomes. The presence of many introns within young megasatellites suggests a model in which an exon-intron DNA segment is first duplicated and amplified before the accumulation of mutations in intronic parts partially erase the tandem repeat in such a way that it becomes detectable only in exonic regions. In addition, evidence for the genetic transfer of megasatellites between unrelated genes suggests that megasatellite formation and evolution is a very dynamic and still ongoing process in vertebrate genomes.

## Introduction

Eukaryotic genomes are characterized by an increase in complexity, often associated with a remarkable expansion of repeated sequences, as compared to prokaryotes (Richard et al. 2005). DNA repeats are traditionally classified in two classes: dispersed repeats and tandem repeats. Among the latter a distinction is usually made between microsatellites, whose basic motif is short (one to nine nucleotide long) and minisatellites, whose basic motif is at least 10 nucleotides long. Although microsatellites have been widely studied in several completely sequenced organisms (Bachtrog et al. 1999; Dieringer and Schlötterer 2003; Hennequin et al. 2001; Innan et al. 1997; Malpertuy et al. 2003; Richard et al. 1999; Röder et al. 1998; Sakamoto et al. 2000; Dib et al. 1996), few publications extensively describe the distribution and variability of minisatellites in eukaryotic genomes. Three independent studies showed that *Saccharomyces cerevisiae* contain 44-111 such repeats, depending on methods and thresholds used for detection (Verstrepen et al. 2005; Richard and Dujon 2006; Bowen et al. 2005). In *Tetraodon nigroviridis*, two minisatellite families were detected, one at the end of subtelocentric chromosomes (10 bp tandem repeat) and the second one at centromeric sequences (118 bp tandem repeat) (Roest Crollius et al. 2000). Previous analyses on individual chromosomes suggest that roughly 3,000-4,000 minisatellites should be found in the complete human genome, mainly located in subtelomeric regions, whereas *Arabidopsis thaliana* and *Caenorhabditis elegans* should contain approximately 720 and 650 minisatellites, respectively (Vergnaud and Denoeud 2008). Unfortunately, no comprehensive work describing minisatellite distribution and variability in several totally sequenced eukaryotic genomes has been published yet.

*Candida glabrata* is an opportunistic pathogenic yeast, responsible for candidiasis and a growing number of nocosomial pathologies, due to its resistance to azole derivatives (Waldin et al. 2008). Its genome was completely sequenced in 2004 (Dujon et al. 2004) and its analysis revealed an unexpected number of large minisatellites, whose basic motif lengths ranged from 126 to 429 nucleotides, covering several kilobases of DNA. These repeats were always found within open reading frames of genes suspected to be involved in cellular adhesion and pathogenicity. They were called “megasatellites” in order to distinguish them from classical minisatellites made of smaller motifs (Thierry et al. 2008, 2009). Further molecular studies using clustering algorithms on *C. glabrata* megasatellite families suggested three independent modes of propagation for these large tandem repeats: i) gene duplication, ii) ectopic gene conversion and iii) transfer of motifs between megasatellites, this last mechanism being tentatively proposed to explain some of the faster evolving motifs (Rolland et al. 2010). More recently, a large survey encompassing 21 fully sequenced fungi genomes confirmed that *C. glabrata* exhibited more megasatellites than any of the other sequenced fungal genomes, at that time. In addition, it was shown that different motifs were found in orthologous genes whereas evolutionary related motifs were found in non-orthologous genes, suggesting that megasatellites are constantly created and lost and that their formation and spreading does not necessarily follow the evolution of their host genes (Tekaia et al. 2013). However, relative mutation rates of fungal megasatellites are difficult to precisely determine due to the lack of appropriate molecular clocks in fungal genomes.

Vertebrates are much younger eukaryotes than fungi. They emerged as a monophyletic group from the Chordate phylum, 550 million years ago, during the pre-Cambrian explosion (Supplemental Figure 1, (Bromham et al. 1998; Erwin et al. 2011)). They spread all over the planet, in almost all ecological niches, from the bottom of the oceans to the highest altitudes, from the north pole to the south pole. They are arguably the most successful of the chordates, with more than 66,000 species described (Genome 10K Community of Scientists 2009). The presence of an internal skeleton that can, in specific circumstances, become partially mineralized, allows for the recovery of fossils. Their precise datation associated with genome sequencing data led to the determination of molecular clocks specific to this clade (Wray et al. 1996). In addition, the function of vertebrate genes and of their cognate proteins have been thoroughly studied, and is still the topic of many investigations, partly due to the fact that our own species belongs to this clade. For all these reasons, it was decided to establish the first set of vertebrate megasatellites, based on the complete sequence of 61 vertebrate genomes, in addition to the yeast *Saccharomyces cerevisiae*, whose mini- and megasatellite content had previously been determined (Richard and Dujon 2006). We detected more than 14,000 tandem repeats whose base motif was longer than 90 nucleotides, unevenly distributed among the 12 clades studied. Two bursts of megasatellite formation were identified, the first one in the gnathostomata, after the agnatha radiation (jawless fish), and the second one in the therians (mammals with uterus). Three quarters of these megasatellites encode zinc-finger proteins, but other cellular functions are highly represented, such as cell membrane, intracellular trafficking and RNA metabolism. Surprisingly, although most megasatellites are encoded in the exonic parts of genes, a significant fraction of them overlap exon-intron junctions in primates, suggesting their recent formation.

## Results

In the present study, a megasatellite was defined as a direct and contiguous tandem repeat of three motifs or more, each of individual length at least 90 bases. We initially planned to search whole genome sequences with two programs: Tandem Repeat Finder (TRF) (Benson 1999) and MREPS (Kolpakov et al. 2003). TRF is a software commonly used to detect tandem repeats, whose algorithm is based on the statistical validation of a tandem repeat by comparison with random sequences (Benson and Waterman 1994).

MREPS uses a combinatorial algorithm considering that two adjacent sequences are part of the same tandem repeat if they differ at most by *k* mismatches. Both software were run on 61 vertebrate species genomes and on the well described *Saccharomyces cerevisiae* genome. These 61 species were chosen because when this work started their whole genome sequence was available and annotated in the *Ensembl* database (release 101) (Yates et al. 2020). They are detailed in Table 1. *Danio rerio* (zebrafish), *Macropus eugenii* (wallaby) and *Tarsius syrichta* (tarsier) genomes contained an abnormally high number of tandem repeats as compared to all other vertebrate genomes, suggesting that sequence quality and/or its assembly were not acceptable to allow faithful detection of real tandem repeats. They were therefore discarded from subsequent studies. The 58 remaining vertebrate genomes contained altogether 4,234 sequences responding to the megasatellite definition. Careful examination of these tandem repeats demonstrated the presence of many false negative as well as false positive results. False positives were typically tandem repeats of transposons, especially SINEs in primate genomes. The human genome, for example, contains more than 1,500,000 *Alu* elements, mostly found in intronic or intergenic regions and they tend to cluster (Richard et al. 2005; International Human Genome Sequencing Consortium 2001). This clustering led to their frequent and wrong identification as tandem repeats by MREPS. False negative corresponded to genes previously known to contain large tandem repeats that were not identified in our analysis, such as WD40 repeats (Hu et al. 2018, 40). Thorough analysis of these false negative revealed that the presence of large introns -frequent in vertebrates- precluded the correct identification of megasatellites encoded in exons. All these results led to the conclusion that directly looking for megasatellites in DNA sequences was not possible to reliably achieve on large and complex vertebrate genomes, with the available tools. We therefore decided to switch to an alternative strategy.

Previous analyses showed that almost all large tandem repeats identified in *S. cerevisiae* genomes, as well as those identified in 20 other fungal genomes, belonged to coding regions (Richard and Dujon 2006; Thierry et al. 2008; Tekaia et al. 2013). We hypothesized that the same was true for vertebrate genomes, or at least that we would identify most -if not all- megasatellites by scanning proteins instead of genomic DNA. Therefore, all protein sequences were extracted from the 58 vertebrate and *S. cerevisiae* genomes. The T-REKS software (Jorda and Kajava 2009) was run on these sequences and the pipeline described in Figure 1 was followed (see Methods for parameters). T-REKS identified 114,918 protein tandem repeats (TR, Figure 1, step 1).

**Figure 1:**
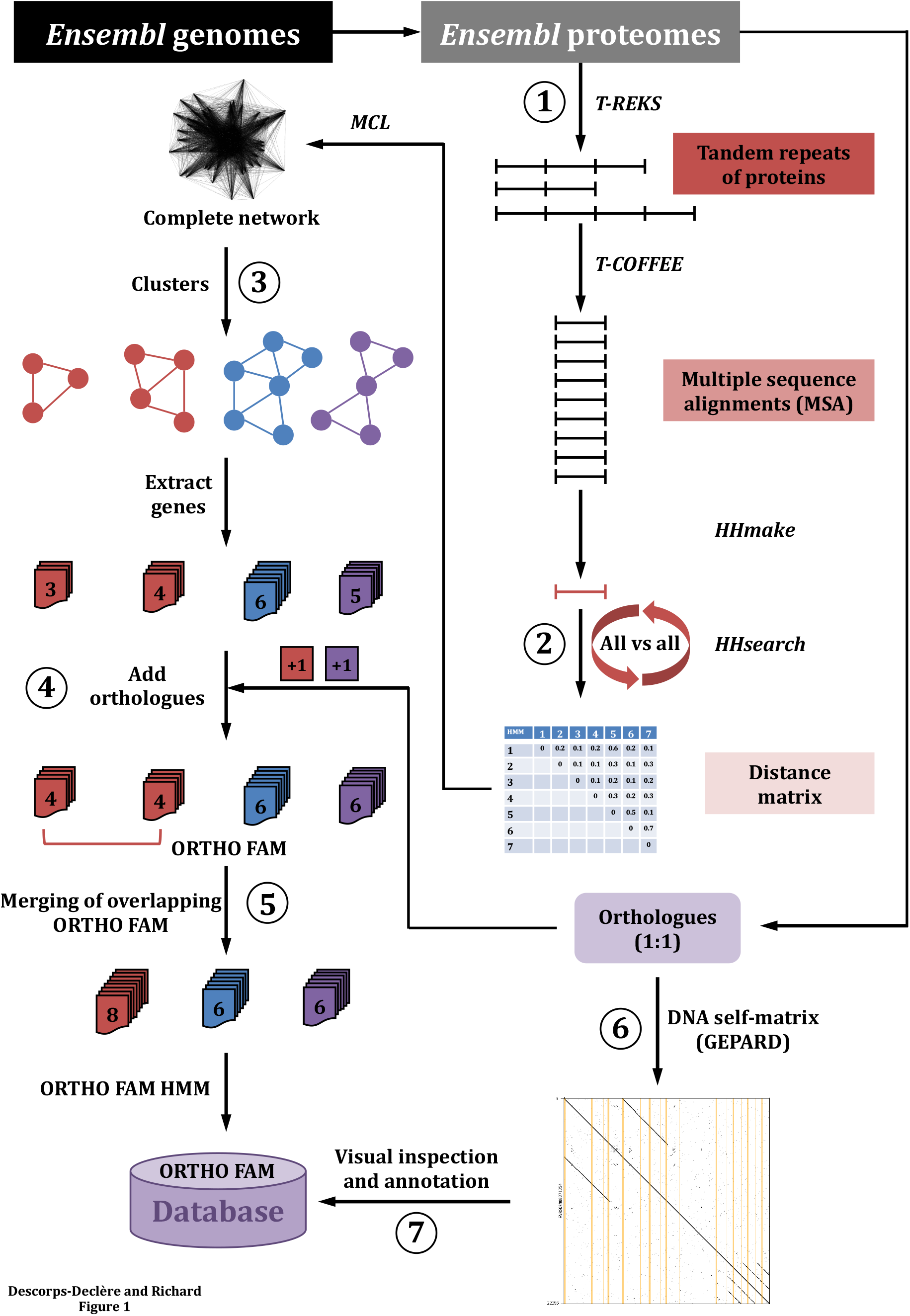
Pipeline used for the analysis. See text for details. Name in italics indicate software used in this analysis.

These repeats exhibit a bimodal distribution, with a first peak showing a very large excess of small motifs (Figure 2).

**Figure 2:**
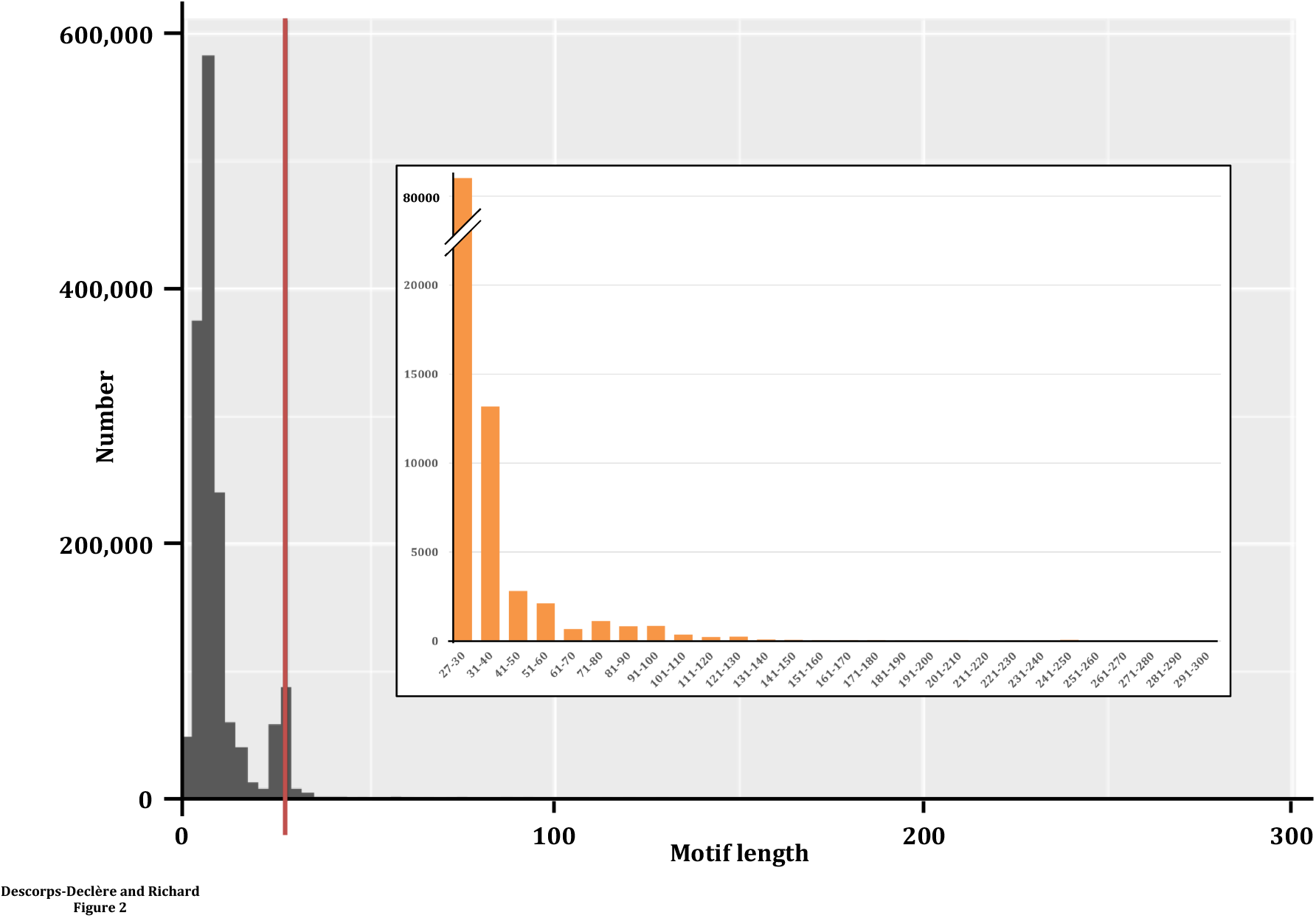
Length distribution of motifs detected by the T-REKS program. A bimodal distribution was observed, separating very short motifs (left of the red line) from longer motifs that were kept for further analyses (right of the red line). Motif lengths are in amino acids. Inset: size distribution of motifs at least 27 amino acids long.

The second peak was centered on a length of 27-30 amino acids, with a large excess of 28 amino acid motifs, corresponding to zinc-finger proteins. The 106,581 TR containing motifs of this length or larger were kept for subsequent analyses (Figure 2, inset). Distances between all Hidden Markov Models were determined to draw a complete network of all distances between HMM (HMM, Figure 1, step 2). From this network, clusters were identified, each corresponding to a TR family (Figure 1, step 3). Genes encoding these TR were identified and all 1:1 orthologous genes in which a megasatellite was found were extracted from *Ensembl* (Figure 1, step 4), and HMM were used to search TR in these orthologues. This step was set up to verify that no TR had been missed by the initial T-REKS search, due to degeneracy of the repeat tract. Depending on how the megasatellite was phased by T-REKS, some TR belonged to the same family, with a different phasing. These were tagged as identical HMM and corresponding ORTHO FAM were merged (Figure 1, step 5).

For each gene in which a megasatellite was identified, transcript annotations were recovered. From these annotations, exon and intron coordinates were extracted and self-DNA matrices were run, for each megasatellite-containing gene (Figure 1, step 6). Each of these 3,982 individual matrices was visually inspected and validated or discarded from the database if no megasatellite was visible (Figure 1, step 7). At this stage, megasatellites were manually annotated as being contained in only one exon (MONOMEGA), more than one exon (MULTIMEGA), or overlapping at least one intron-exon junction (OVERMEGA). In some cases, the megasatellite was spread over several small exons separated by large intronic regions. These were called HIDDEN MULTIMEGA in the database, but were considered as MULTIMEGA in all subsequent analyses. Altogether, 3982 megasatellites were identified, distributed among 142 families including at least one member in one species. To this number must be added 10,575 zinc-finger proteins that were treated separately (see below). These families were called ORTHO FAM, the majority of them being either MONOMEGA (65), MULTIMEGA (46) or OVERMEGA (26), while four contained a mix of more than one megasatellite type and were annotated as MIXMEGA (Supplemental Table S1). Finally, one megasatellite was inadvertedly found in an intronic sequence, and conserved in 27 eutherian species. This suggests that either it plays a functional role or that this gene annotation is wrong and this megasatellite is not purely intronic. Some examples of each category are shown in figure 3. Since whole-genome duplications were frequent during vertebrate evolution (Dehal and Boore 2005; Jaillon et al. 2004), only these 142 ORTHO FAM families were considered for further analyses and megasatellites encoded within paralogues were discarded at this stage.

**Figure 3:**
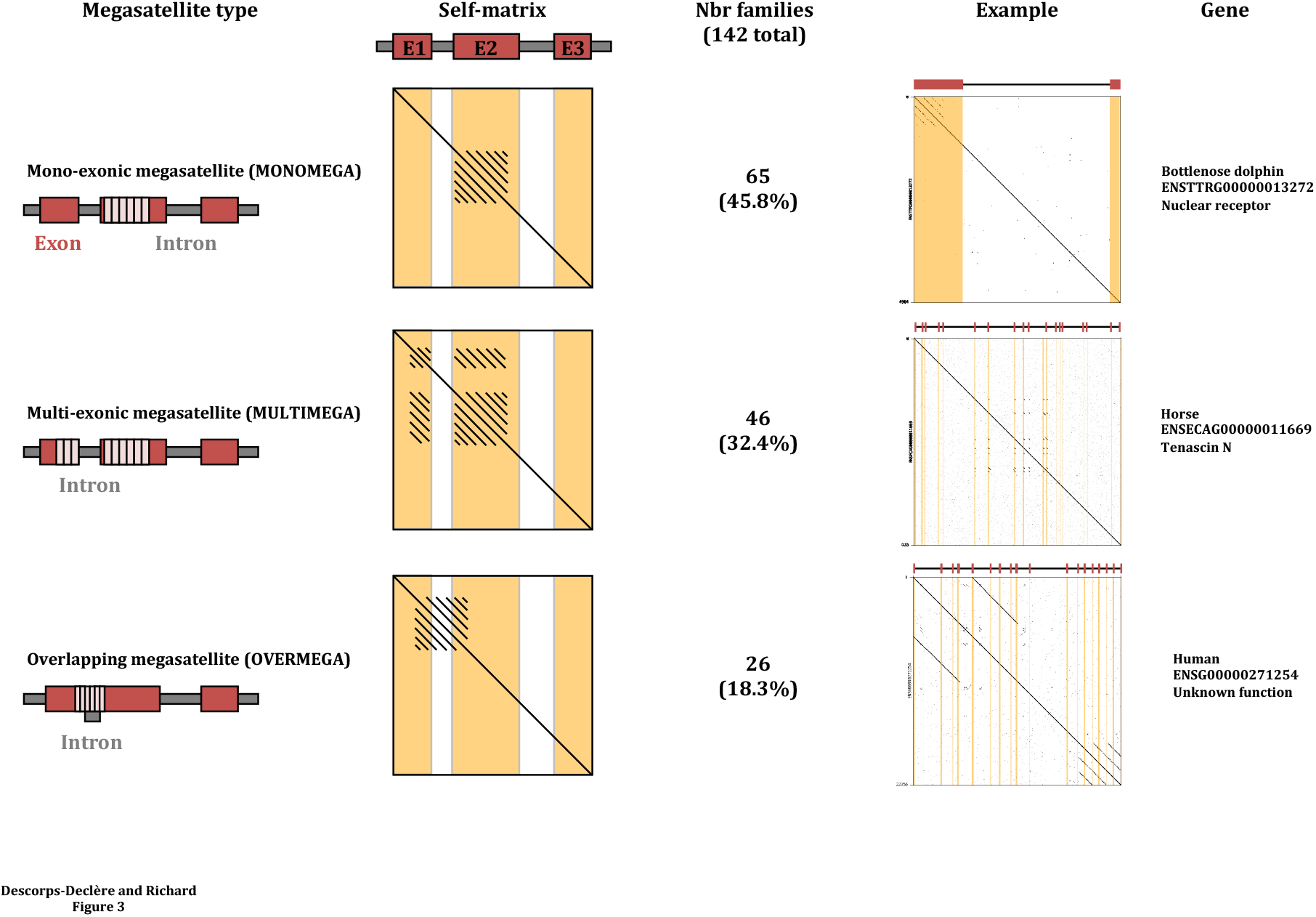
Megasatellite types. **Left**: Monomega, multimega and overmegasatellites are sketched. Red bars: exons. Grey bars: introns. **Middle**: Schematic representation of self matrix dot plots, for each type. **Right**: One example of each megasatellite type is shown. The longest transcript is shown above each dot plot.

### Distribution of megasatellites among vertebrate clades

All vertebrate clades were found to contain megasatellites, although to different extents. Primates and eutherians contain respectively 116 and 107 out of 142 families, whereas only 17 families were detected in agnatha (Supplementary Table S2). Eight of them are common to all vertebrate clades, Tenascin C (ORTHO FAM 1130) and Tenascin R (ORTHO FAM 4068), two related developmental proteins, Cortactin (ORTHO FAM 1163), an actin polymerization protein, growth factor beta binding protein 1 (ORTHO FAM 1746), Nebulin (ORTHO FAM 108) and Nebulette (ORTHO FAM 175), two proteins essential to regulate the stability and length of actin filaments in skeletal and cardiac muscle fibers, CREB1 (ORTHO FAM 113), involved in cAMP response and Angiomotin (ORTHO FAM 4295) involved in cell motility (Figure 4A). Five families are present in all gnathostomata (jawed vertebrates), but two of them were lost in monotremes (Figures 4B and C). Seven families were common to all sauropsida and mammals, except monotremes (Figure 4D). It is however possible that due to uncomplete assembly of the platypus genome sequence (the only monotreme sequence available) these megasatellites may have been missed in this species (Warren et al. 2008).

**Figure 4:**
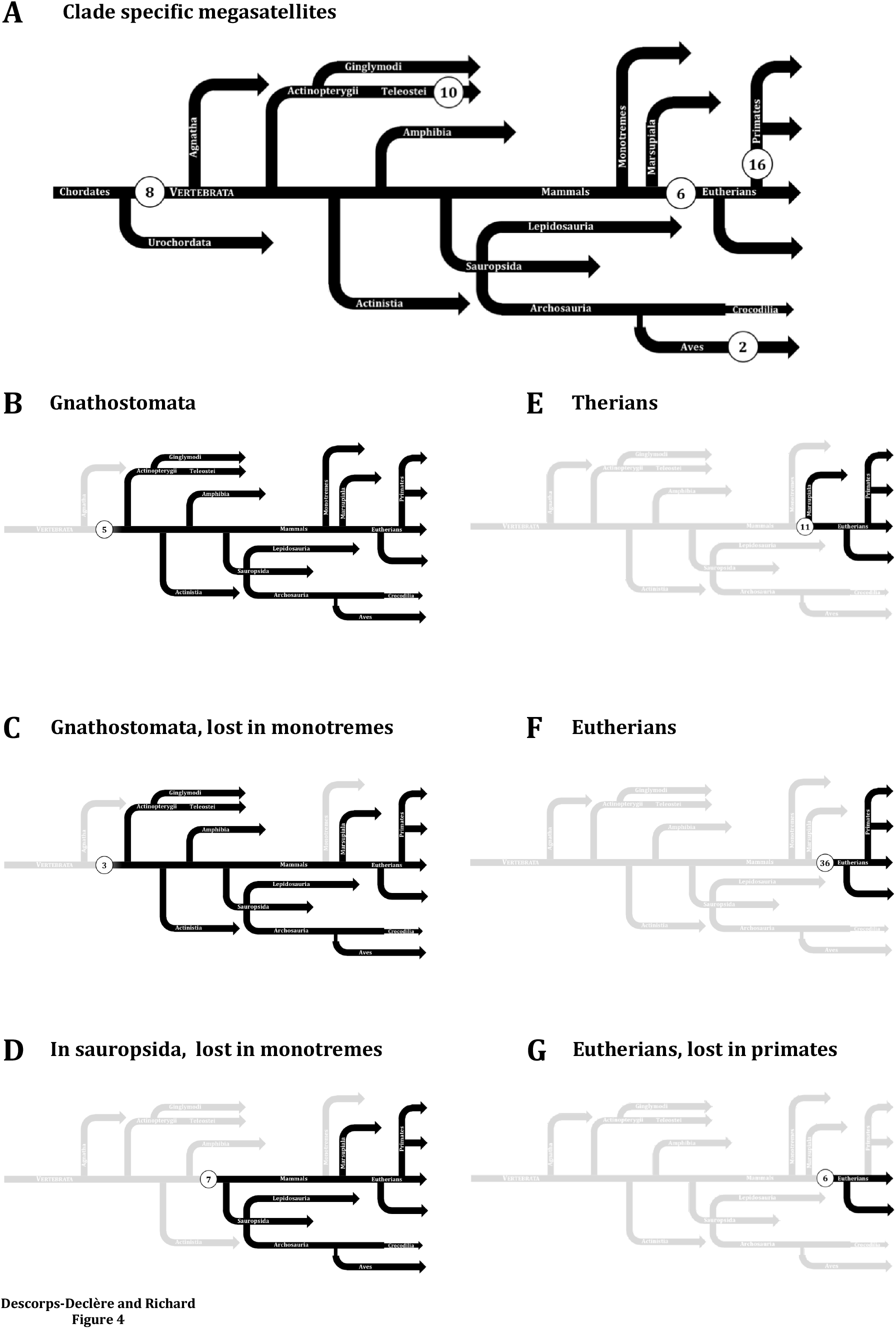
Megasatellite distribution in vertebrate clades. The number of clade-specific megasatellites are shown in **A**. Other part of the figure (**B-G**) identify megasatellites that are common to one or more vertebrate clades, as indicated.

On the opposite, clade-specific families seem to be frequent in fishes and primates. The latter contains 16 megasatellite families that are absent from other clades, whereas teleostei contain 10 specific families (Figure 4A). However, the number of species studied, hence of proteomes varies among clades. When correction was made for proteome sizes, it was found that agnatha, amphibia, sauropsida and monotreme exhibited significant less families than other clades (Chi2 p-value<0.01). In addition, therian genomes (marsupials, eutherians and primates) contain significantly more families than any other genome studied (Chi2 p-value<0.01). We concluded that there were probably two bursts of megasatellite formation during vertebrate evolution, one at the root of the clade, after the agnatha radiation and another one much larger in mammals, after the monotreme radiation (Figure 5).

**Figure 5:**
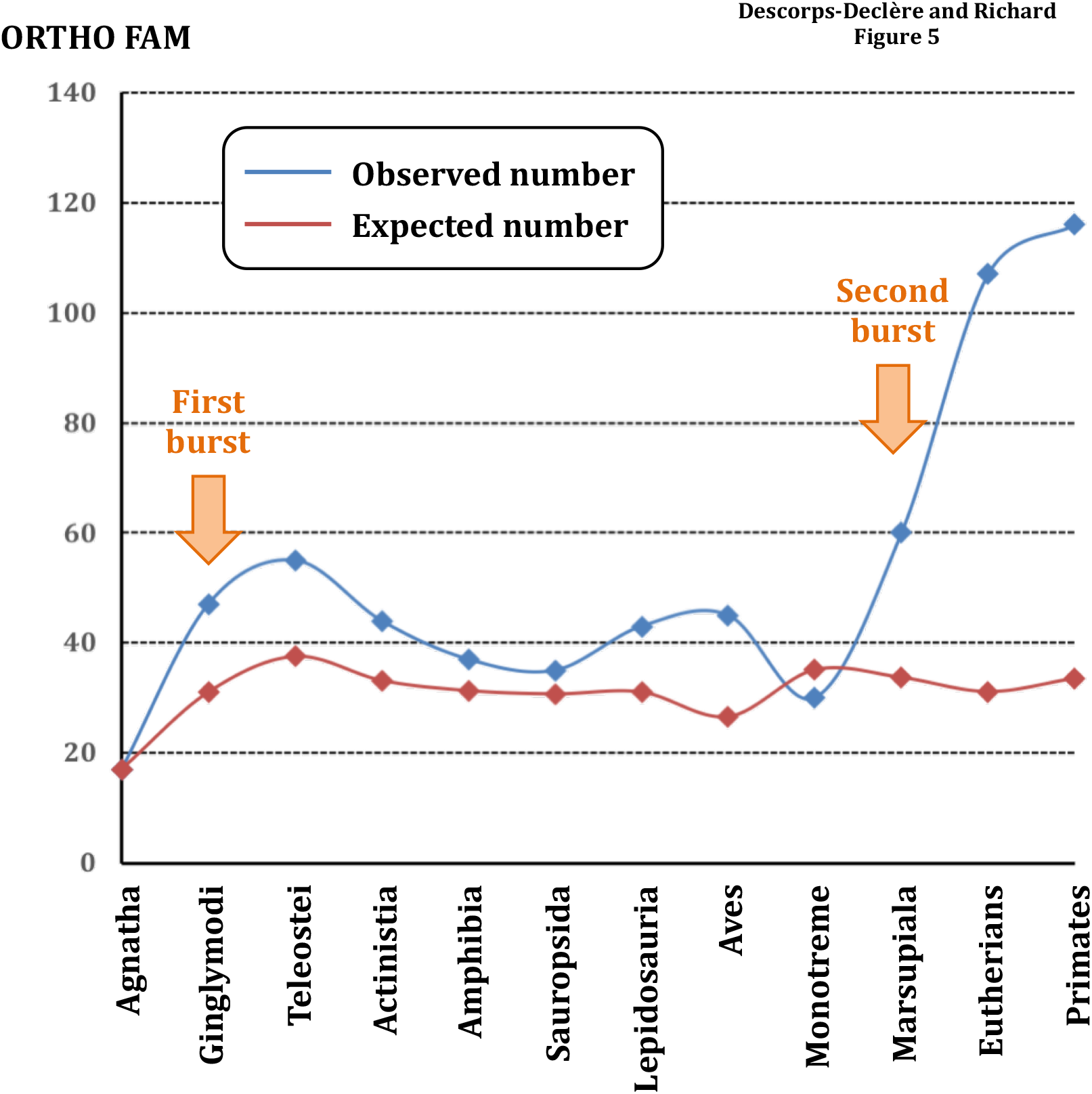
Number of expected versus observed number of megasatellites in each clade. The two orange arrows point to the two statistical increases in megasatellite number during vertebrate evolution.

It is unclear whether the significant decrease of families in amphibia, sauropsida and monotreme reflects a biological reality, technical limitations in sequencing and/or assembly of these genomes or the rather limited number of species sequenced (only one in each if these clades). Finally, 11 families were common to therian mammals (4E), 36 families were found to be conserved in all eutherians (Figure 4F) while six were specific of non-primate eutherians, like mouse, dolphin and bats (Figure 4G).

In order to check whether megasatellite distribution was homogeneous within each clade, their occurences were compared in clades containing more than one species, i.e. birds, marsupiala, teleosteans fishes, eutherians and primates.

A significant excess of megasatellites was found in two birds (*Ficedula albicollis* and *Meleagris gallopavo*), two fishes (*Oryzias latipes* and *Poecilia formosa*), and two mammals (*Equus caballus* and *Mus musculus*) (Chi2 p-value<0.01). Besides these few cases, the number of megasatellites in each species was very homogeneous within a given clade.

### Distribution and function of megasatellites among species

Altogether, 3,982 megasatellites were detected in the 59 yeast and vertebrate families, not including zinc-finger protein-encoding genes (see below). Not surprisingly, the species with the fewest repeats is *Petromyzon marinus* (19 megasatellites) while *Mus musculus* has the highest number of megasatellites (133, Table 1). Since paralogous genes were discarded during the analysis, only one member of each ORTHO FAM family was represented in each species, hence the number of megasatellites per species is equal to the number of families present in that species.

Megasatellite-encoding genes were classified according to their known (or inferred) function. The most frequent ones were linked to cell membrane metabolism (15%), intracellular trafficking, including interactions with actin, myosin and tubulin (11%) and RNA metabolism (10%). A significant fraction of them (8%) could not be associated with any known function, based on sequence homology or protein motifs (Figure 6A). Most of the megasatellites (68%) encoded in the eight more frequent functional categories are present in all clades, with the exception of those involved in immune response that are absent from sauropsida and monotreme (Figure 6B).

**Figure 6:**
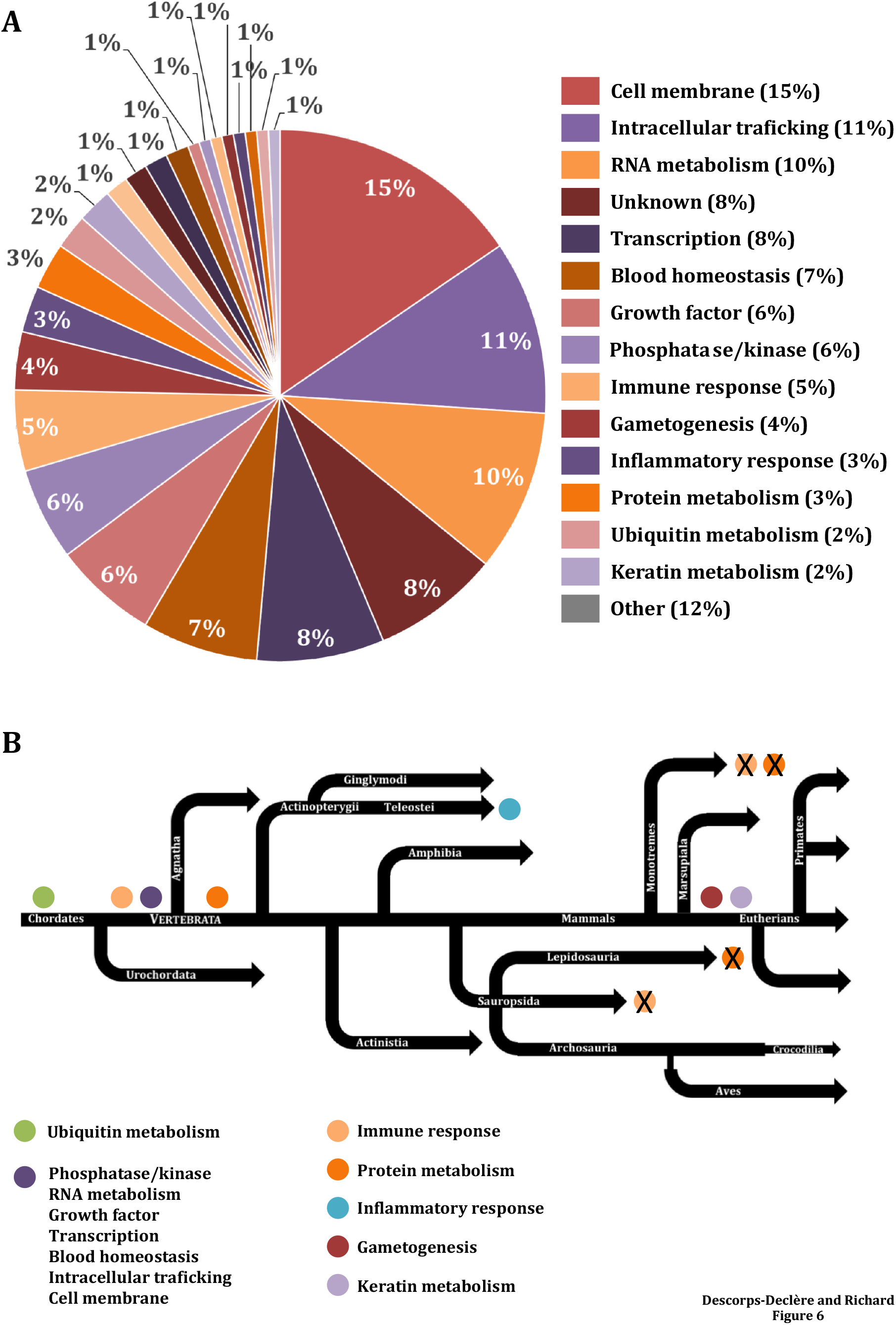
Function of megasatellite-encoding genes. **A**: Pie chart showing the frequency of all identified functions. “Other” encompasses all functions associated with 1% or less genes. **B**: Repartition of most frequent functions along the vertebrate tree. Presence of a function is indicated by a colored disk, absence by a crossed colored disk.

Megasatellites in genes playing a function in protein metabolism are present only in gnathostomata, to the exception of lepidosauria and monotreme. Several megasatellites are found in genes dedicated to a specific function and are unique to certain clades. It is the case of those related to the inflammatory response that are detected only in teleostei, and those involved in gametogenesis or keratin metabolism that are found only in eutherians (Figure 6B). These results show that, to a few exceptions, most megasatellites are encoded in genes that are common to all vertebrate clades and are therefore involved in conserved functions.

### Megasatellites propagate by two different mechanisms

In the course of the present study, several orthologous families were identified as containing more than one member per species. This is normally impossible, since only 1:1 orthologous genes in which at least one member contained a megasatellite were kept. A more thorough analysis of these 24 families showed that they actually included paralogous genes. If both paralogues carry the same megasatellite, independently identified at the beginning of the pipeline (Figure 1, step 1), they ended up in the same family after clustering (Figure 1, step 3).

These families showed that they could be classified in three different cases. Besides the 10 families whose sequence quality was too low to obtain reliable alignments, the simplest case included real paralogues containing the same megasatellite at the same position, in each species. This was the status of six families (ORTHO FAM 143, 156, 1163, 3045, 3121 and 3340). An example of such a paralogous megasatellite is shown in figure 7A.

**Figure 7:**
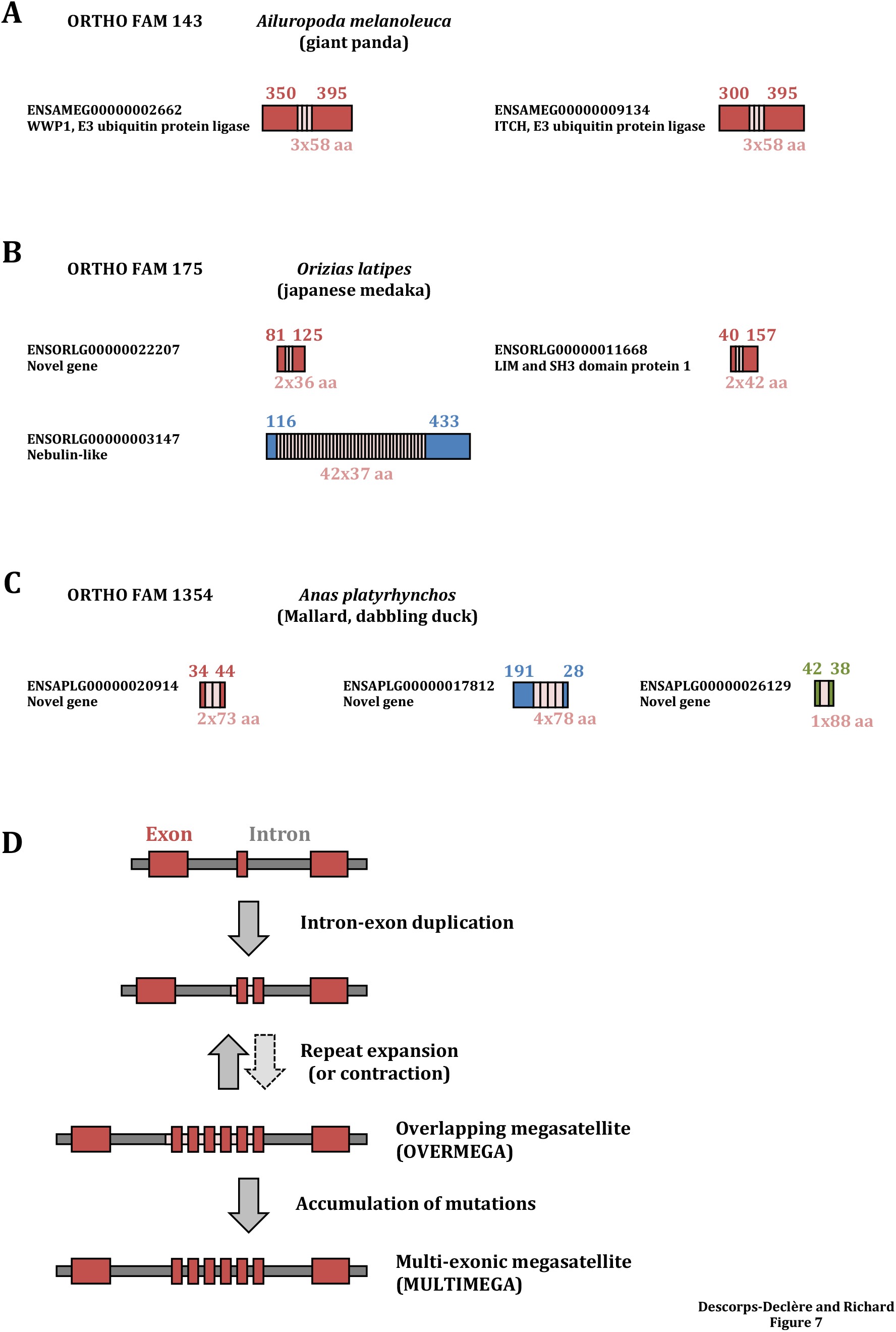
**A**: An example of two paralogous genes containing the same megasatellite. The family as well as the species are indicated above the two paralogues. Gene names and supposed functions are shown next to the protein. Lengths of each protein part are indicated in amino acids. **B**: An example of a three-member family containing two paralogues. Homologous protein parts are in red, the third protein is in blue. The megasatellite is present as a duplication in the two paralogues and is largely expanded in the third member. **C**: An example of a three-member family containing three unrelated genes (in red, blue and green). The megasatellite is present as a single motif in the green gene. **D**: A model for megasatellite evolution in vertebrates. See text for details.

More interestingly, in four families (ORTHO FAM 113, 175, 1130 and 3163) more than two genes containing the same megasatellite were identified. Two of these genes are paralogues, whereas the third one is unrelated. This suggests that the megasatellite was transferred, or jumped, from one gene to another one, although we cannot completely exclude that the third gene is a paralogue that has diverged from the two others more rapidly than its megasatellite (Figure 7B). A third case corresponds to only one small family (ORTHO FAM 1354) in which the same megasatellite is found in three different genes in one species. However, the megasatellite sequence is not totally conserved and its length differs among the three genes, one of them containing only one motif, unrepeated (Figure 7C). Finally, three families contained short proteins, Ubiquitin C (UBC, ORTHO FAM 3898) and two others too small to detect a reliable homology (ORTHO FAM 1211 and 1864) between their non-repeated parts. We concluded that megasatellites appear to propagate by two different modes, one involving duplication of an existing megasatellite-containing gene and another unexpected one, relying on transfer from one gene to another, phylogenetically unrelated, gene.

### Megasatellite length does not increase with time

Ubiquitin is a eukaryote-specific gene of archaeal origin, encoding a tandemly repeated 76-amino acid polypeptide. It is post-translationally cleaved into the active 76-residue ubiquitin peptide, involved in regulating protein metabolism (Grau-Bové et al. 2015). In budding yeast, *UBI4* was found to exhibit variability in the number of 76-amino acid tandem repeat units among different strains (Gemayel et al. 2017). In several eukaryotic lineages, *UBI4* is duplicated, as two paralogous genes called UBB and UBC, the latter exhibiting more repeat units than the former. Using the present approach, we have detected polyubiquitin in 47 out of 59 species studied (ORTHO FAM 3898). In some cases, two genes were identified as containing a polyubiquitin repeat. In these cases, it was assumed that the two copies correspond to UBB and UBC. When only one gene was detected, we assumed it was UBC, UBB being the shorter one (Gemayel et al. 2017). The number of repeat units identified is generally in good accordance with previous reports. The average number of repeats is 5 but a large size variation is observed around this mean value (from 2 to 20 repeat units). There is no visible expansion of the ubiquitin megasatellite length during vertebrate evolution. However, each clade exhibits a large variability among species (Supplemental Table S3).

When all other megasatellite families were considered, the average number of motifs per tandem repeat was remarkably constant, around 7 per megasatellite in most clades, ranging from 6.3 in ginglymodi to 10.9 in agnatha (mean= 7.7, 99% c.i.= 6.4-9.0). This indicates that megasatellites do not tend to increase in length with time.

### Abundance of zinc-finger protein families in primates and lepidosauria

Zinc-finger proteins (ZFP) are encoded by tandem repeats that were identified as megasatellites in the present study. Altogether, 10,575 ZFP were found, unevenly distributed in all clades. There is a general increase in ZFP number with time, younger vertebrates, encoding more such megasatellites than older clades (Figure 8).

**Figure 8:**
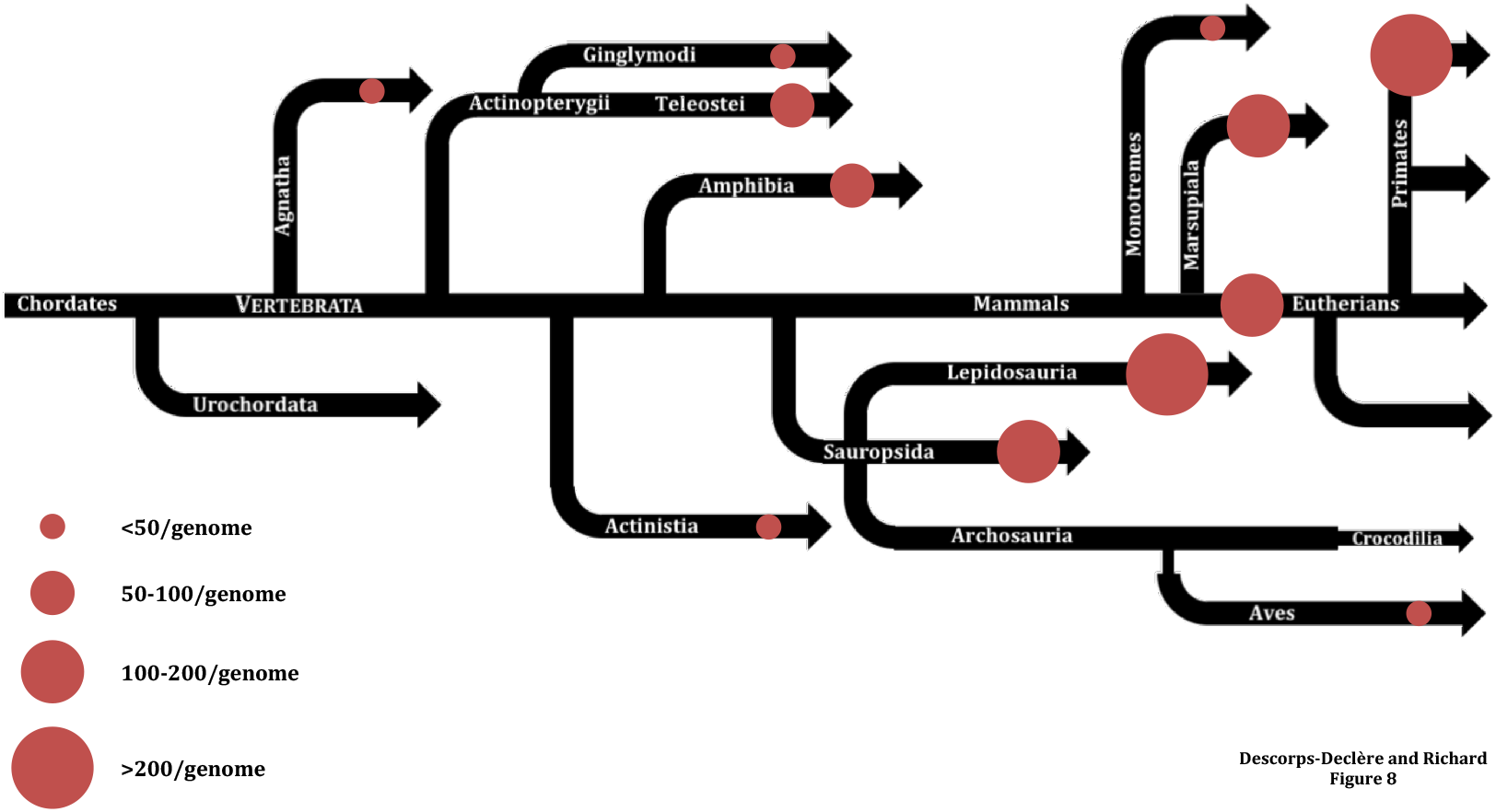
Zinc-finger gene frequency in vertebrate genomes. The size of the colored disk represents the average number of megasatellites in each genome.

When proteome size of each clade was taken into consideration, ZFP number was significantly higher in sauropsida, lepidosauria, marsupiala, eutherians and primates (Fisher exact test p-value<2.2 x 10^-16^). Comparisons between these five clades showed that they were more frequent in primates and in lepidosauria than in the three others and more frequent in lepidosauria than in primates (Fisher exact test p-value= 0.002). We concluded that ZFP megasatellites were expanded in lepidosauria and more frequent than in any other vertebrate clade.

Krüppel-associated box domain-containing zinc-finger proteins (KZFP) are a subclass of ZFP that regulate transposable elements by repressing their expression. We found that KZFP were restricted to four families, ORTHO FAM 1, 3191, 4452 and 5539, the three last families containing only five KZFP. They are less frequent in aves than in any other vertebrate clade, confirming a former study (Imbeault et al. 2017).

The 10,575 ZFP megasatellites are distributed among 464 different families, one of them (ORTHO FAM 1) containing 86% of them. It must be noted that ORTHO FAM 1 contains the highest number of ZFP megasatellites in all clades, except for lepidosauria, in which it is another family (ORTHO FAM 1033) that is the most preeminent. Careful examination of these two families showed that they actually contained the same megasatellite that was phased differently (Supplemental Figure 2). This issue was normally addressed in the analysis pipeline (Figure 1, step 5), and these two families should have been merged, like other overlapping ORTHO FAM. However, like hereabove explained the presence of paralogues within each of these two families prevented them to be merged during the analysis.

## Discussion

In the present study, we have identified 14,557 megasatellites encoded in 58 vertebrate and budding yeast genomes (Supplemental Table S4). It is, by far, the most complete description to date of large tandem repeats in eukaryotic genomes. It is remarkable that 18% of non-ZFP megasatellites are overlapping several introns (Figure 3), these cases being more frequent in primates, which are the youngest species studied here, having diverged from other eutherians less than 10 million years ago (Supplemental Figure 1). This is less frequent in older vertebrates, suggesting that mutations accumulated over time in intronic sequences, slowly erasing any trace of tandem repeats encoded within introns. This suggests that megasatellite formation in vertebrates may start with the duplication of an exon-intron DNA segment, that becomes subsequently amplified to form an overlapping megasatellite (OVERMEGA). Mutations accumulate over time within the intronic part, erasing the tandem repeat and leading to what is detected as a multi-exonic megasatellite (Figure 7D). Introns are thought to have invaded the proto-eukaryote following the endosymbiosis of an α-proteobacteria with an archaea ancestor. In support of this hypothesis, group II self-splicing introns show structural and catalytic similarities with spliceosomal introns and α-proteobacteria contain a large number of such elements, as compared to other bacteria (reviewed in (Rogozin et al. 2012)). Intron size tend to increase over time during eukaryotic evolution, ancestral eukaryotes exhibiting smaller introns than more recent ones. (Csuros et al. 2011). If a tandem repeat is found overlapping one or more introns (OVERMEGA), two mutational forces will tend to erase its presence over time: i) intron size increase, by accumulation of transposons or other virus-like elements, that will make megasatellite detection impossible for current software; ii) accumulation of mutations that will slowly erase the repeat in the intronic part of the OVERMEGA to transform it into a MULTIMEGA.

### Megasatellites exhibit two modes of propagation

In a former study on *S. cerevisiae* and *C. glabrata* megasatellites, it was shown that these tandem repeats propagate by three different mechanisms: i) duplication of an existing megasatellite-containing gene; ii) megasatellite “jump” between two unrelated genes; iii) gene conversion between paralogues (Rolland et al. 2010). Here, we show that at least two of these mechanisms are recapitulated in vertebrates. We found six families in which all species contained two paralogues encoding the same megasatellite (Figure 7A). These paralogous genes may come from whole-genome duplications, common in vertebrate evolution (Jaillon et al. 2004; Ohno 1970; Sacerdot et al. 2018), or from segmental duplications or Copy Number Variations (Sudmant et al. 2013; Zarrei et al. 2015). The second mechanism implies that a given megasatellite can transfer its genetic information from one gene to another unrelated gene, like a transposable element. Several such examples have been discovered in the present study (Figure 7B and 7C). This strongly suggests that some megasatellites have the propensity to jump from one gene to another one, maybe by using an RNA intermediate followed by reverse transcription and integration or by an alternative yet unknown mechanism. Such integrations must be in frame with the targeted gene, else they would inactivate it.

### Megasatellites are frequently found in cell membrane genes

In yeast and fungi, megasatellite-encoding genes are most of the time involved in cell wall metabolism and function. In *S. cerevisiae*, it is the case of the *FLO* gene family important for cell-cell adhesion and flocculation (Verstrepen et al. 2005; Smukalla et al. 2008; Fidalgo et al. 2006, 11). The large ALS gene family also contains a megasatellite whose role in adhesion has been demonstrated in *Candida albicans* (Oh et al. 2005). In *Candida glabrata*, the megasatellite-containing EPA gene family is also essential for cellular adhesion to human epithelial cells (Cormack et al. 1999). Genome wide identification of large tandem repeats in 21 fungi genomes showed that half of them were encoded in genes expressing plasma membrane or cell wall proteins, involved directly or indirectly in cell-cell adhesion (Tekaia et al. 2013). Here, we show that the most frequent function associated with megasatellite-encoding genes in vertebrates is cell membrane metabolism (Figure 6). This raises the question of a universal role of long tandem repeats in cell membrane homeostasis. The repeated peptide could serve as a linker sequence between the protein part imbedded in the membrane and the part located at cell surface. Increasing or decreasing linker length could modulate interactions between membranes of different cells.

### Microsatellites and megasatellites show strikingly different evolution patterns

Many former studies aimed at characterizing microsatellite distribution in eukaryotes. They generally concluded that their frequency per megabase of DNA greatly varied among organisms, without any visible trend associated with genome complexification (Richard et al. 2008). When microsatellites of different but closely related yeast species were analyzed, it was found that some dinucleotide or trinucleotide repeats that were frequent in one species were underrepresented in another one, independently of overall genome composition (Malpertuy et al. 2003). Hence, the rules underlying *de novo* microsatellite formation are far to be understood, although there is a statistically significant association of *Alu* elements with microsatellites, suggesting that such transposons help to propagate microsatellites in primate genomes (Jurka and Pethiyagoda 1995; Nadir et al. 1996). A more recent study was performed on 71 vertebrate genomes, including all the genomes of the present study. The authors concluded that the *Xenopus laevis* genome contained less microsatellites per megabase of DNA than the other genomes, whereas squamate reptile genomes were comparatively enriched in microsatellites. This microsatellite expansion may be linked to the propagation of LINE elements in this lineage (Adams et al. 2016), reminiscent of what was already observed in primates with *Alu* retrotransposons. In conclusion, microsatellites are more frequent in some vertebrate clades, independently of their evolutionary age or genome size or complexity, whereas megasatellites underwent two bursts of expansion, one after the agnatha-gnathostomata radiation, and another one in the mammalian clade, after the monotreme radiation.

### The next frontier

The *S. cerevisiae* genome contains five megasatellites encoded in *UBI4, FLO1, FLO5, FLO9* (which are three paralogues) and *NUM1* (Verstrepen et al. 2005; Richard and Dujon 2006). With the present approach only *UBI4*, encoding ubiquitin, was identified. This is due to a known limitation of our approach. In order to study megasatellite evolution, it was decided to build families of orthologous megasatellite-containing genes. *NUM1* and *FLO* megasatellites were detected by T-REKS but since they have no orthologues in any vertebrate, they were discarded from the analysis. This limitation could be alleviated in the future by considering all megasatellites, not only those present in more than one genome.

A recurrent limitation of all studies on tandem repeats is sequence length and quality. Vertebrate genomes analyzed here were sequenced using either first generation Sanger sequencing or second-generation short read (Illumina) sequences. In both cases, read length is not sufficient to encompass a whole megasatellite. Therefore, read assembly is an obligate step to reconstitute the complete sequence, with all caveats linked to tandem repeat assembly. For this reason, three genomes that did not seem to comply with high quality sequence standards for repeats were discarded from the present analysis (Table 1). Nevertheless, it is probable that megasatellite length was often underestimated here. Third generation sequencing, using Nanopore or PacBio long reads should help overcoming this problem, when sequence quality of such reads will be improved to the level of alternative technologies. Another issue is allelic polymorphism, since all vertebrate genomes are diploid but only one set of chromosome sequence was available in the database. This issue is of course not specific of tandem repeat analyses and will be improved in the coming years with more efficient and thorough sequence analysis and storage.

Finally, tandem repeat detection is another limitation, since it is known that no algorithm is able to correctly detect all tandem repeats of a given genome (Leclercq et al. 2007). For this reason, we focused the present work on megasatellites found in protein-coding genes, but non-coding regions also contain large tandem repeats, like Copy Number Variations (Sudmant et al. 2013; Zarrei et al. 2015). Progress should therefore also be made in this area, using more intelligent software based on alternative detection methods and machine learning algorithms. This is probably the next frontier of tandem repeat identification in complex genomes.

## Methods

We developed an original method able to capture the genomic structure of orthologous Tandem Repeat (TR)-bearing protein families. As it is difficult to detect megasatellites directly in vertebrate genomes, our analysis consisted of searching for TR-rich orthologous protein families and studying the genomic structures of the genes encoding the members of these families using dotplots. The whole pipeline of analysis is schematized in figure 1.

### Detection of TR-bearing proteins

We executed T-REKS (Jorda and Kajava 2009) on all proteins of all proteomes of our dataset. To keep only the most relevant results, length distribution of all detected tandem repeats was studied. TR length distribution bimodality led us to set a minimum motif length to be superior or equal to 27 amino acid (Figure 2). The number of motifs composing the TR was set to be at least three.

### TR modelling

To capture inter-motif sequence diversity, we formalized protein TR as a Hidden Markov Model (HMM) profile (Anisimova et al. 2015). To do this, each tandem repeat was subdivided into its individual motifs and their sequences were saved in a multi-sequence file. Those files were then aligned with T-COFFEE (Notredame et al. 2000) and resulting Multi Sequence Alignments (MSA) were filtered to retain only columns with less than 50 % gaps.

### TR clustering

Distances between MSA were calculated with HH-suite (Söding 2005). In order to group related TR (e.g. belonging to orthologous proteins), we instantiated a TR similarity graph from which clusters of strongly similar TR were extracted using the MCL method (van Dongen and Abreu-Goodger 2012). Practically, we first transformed each MSA into an HMM using the hhmake tool, using default parameters. Then we concatenated all produced HMM profiles into a giant HMM dataset. In order to generate similarity matrix of all TR, each profile was compared to all others, running a HHsearch against the full dataset. All results were then transcoded from hhr formatted results to “abc” format (https://micans.org/mcl/man/clmprotocols.html). The E-value was retained as the distance between two HMM, and will be used to draw a complete similarity graph using MCL, which implements a fast and scalable unsupervised cluster algorithm. It was run with Inflation parameters set to 1.4 in order to cut the graph into clusters, each of them representing ancestrally related TR.

### Constitution of ORTHO FAM families

From each MCL cluster, TR-carrying proteins composing this cluster were extracted. For each protein, the API Rest of Ensembl (Yates et al. 2020) was used to retrieve orthologues in the 58 vertebrate genomes eventually kept. These orthologues were added to the protein lists to make protein families. These consolidated families, called ORTHO FAM, are composed of TR-bearing proteins and of orthologous proteins that may or may not contain a TR.

### Merging of differently-phased TR

This step addresses the problem of TR phasing. Depending on the first amino acid used to define the TR, motif sequences may not be identical in the end. For example, the sequence ABCD ABCD ABCD may be phased as a TR whose motif is ABCD, or BCDA, CDAB, or DABC. Therefore, they could possibly be gathered in different TR clusters. To circumvent this problem, a script producing an ORTHO_FAM inclusion graph was developed. Each node is a protein family and each edge symbolizes a shared protein. Using this graph, families containing strongly connected components (homologous proteins) were merged. This allowed to merge TR-containing proteins which were differently phased by T-REKS.

### Database and dotplots

These new ORTHO FAM were deposited in a SQLite database. For each protein component of these families, genomic sequence and gene model corresponding to each protein were retrieved. Using this information and a modified version of Gepard (Krumsiek et al. 2007), 7,533 dotplots for all the genes encoding these proteins were generated. Families containing only one gene were discarded. The remaining 3,982 dot plots were all visually inspected and manually annotated as MONOMEGA, OVERMEGA, MULTIMEGA, HIDDEN MULTIMEGA, MIXMEGA or INTROMEGA. Each of these annotations corresponds to an identifiable pattern on the dotplots, as illustrated for MONOMEGA, OVERMEGA and MULTIMEGA in Figure 3. All megasatellites are shown in Supplemental Figure S4 and all dot plots are in Supplemental Material S1.

## Data access

Scripts used for the analysis were deposited on Github (sdeclere/tandem_repeats). The analysis pipeline is described in pipeline.md.

## Ackowledgements

Work in G.-F. Richard laboratory is generously supported by the Institut Pasteur and by the Centre National de la Recherche Scientifique (CNRS).

## Author contributions

SDD and GFR designed the analysis. SDD wrote the pipeline, set up the database and extracted the results. SDD and GFR analyzed the results and wrote the manuscript.

## Supplemental Figures

**Supplemental Figure 1:**
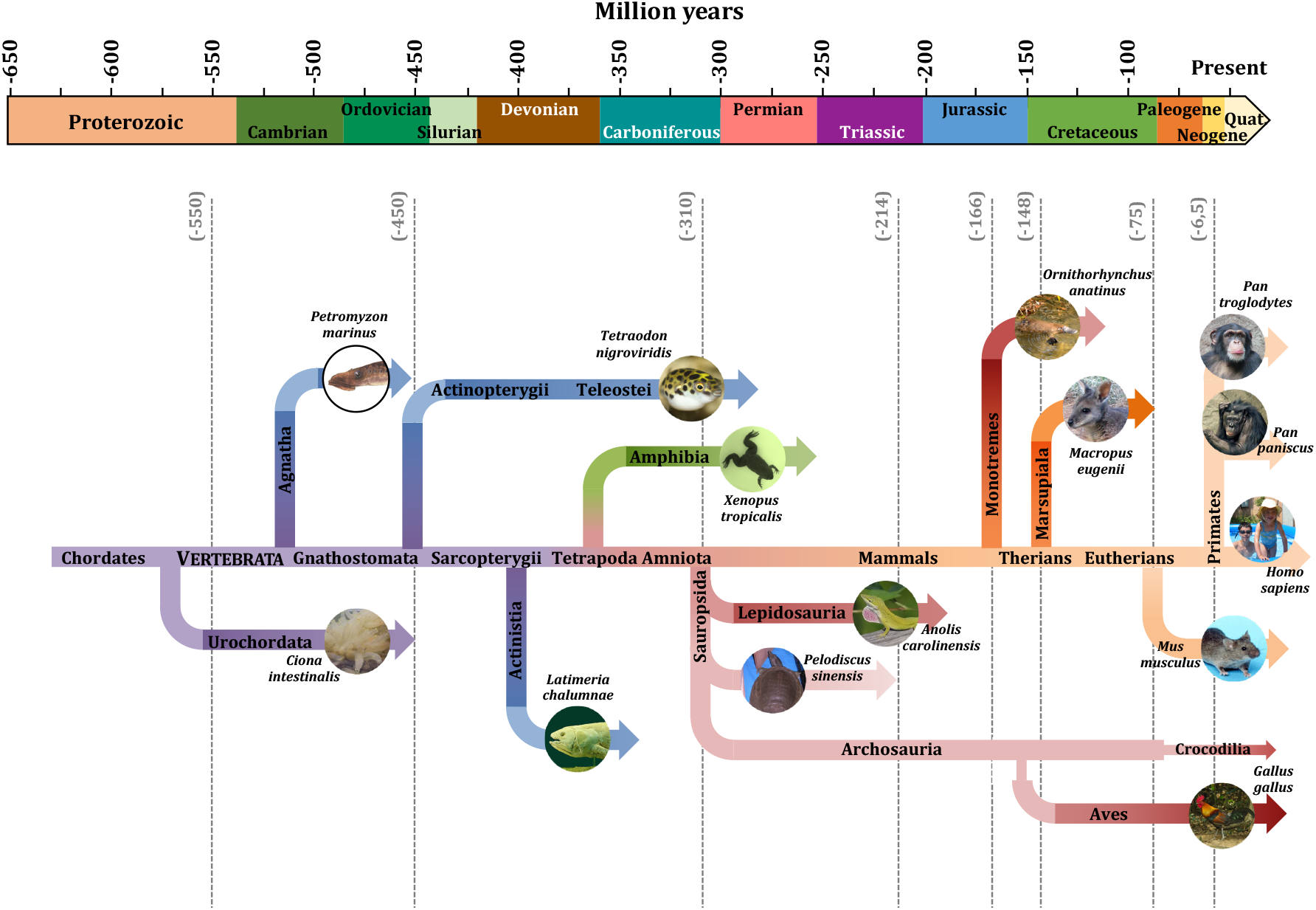
A tree of vertebrate species. Geological periods are indicated at the top, along with time in million years before present. Main vertebrate radiations are indicated, as well as some of the species analyzed here. The constriction between archosauria and aves represents the archaeopteryx, the ancestor of all modern birds. Crocodilians are the only non-flying modern survivors of ancient dinosaurs. Vertical dotted grey lines indicate main radiation dates, as calculated from molecular clocks.

**Supplemental Figure 2:**
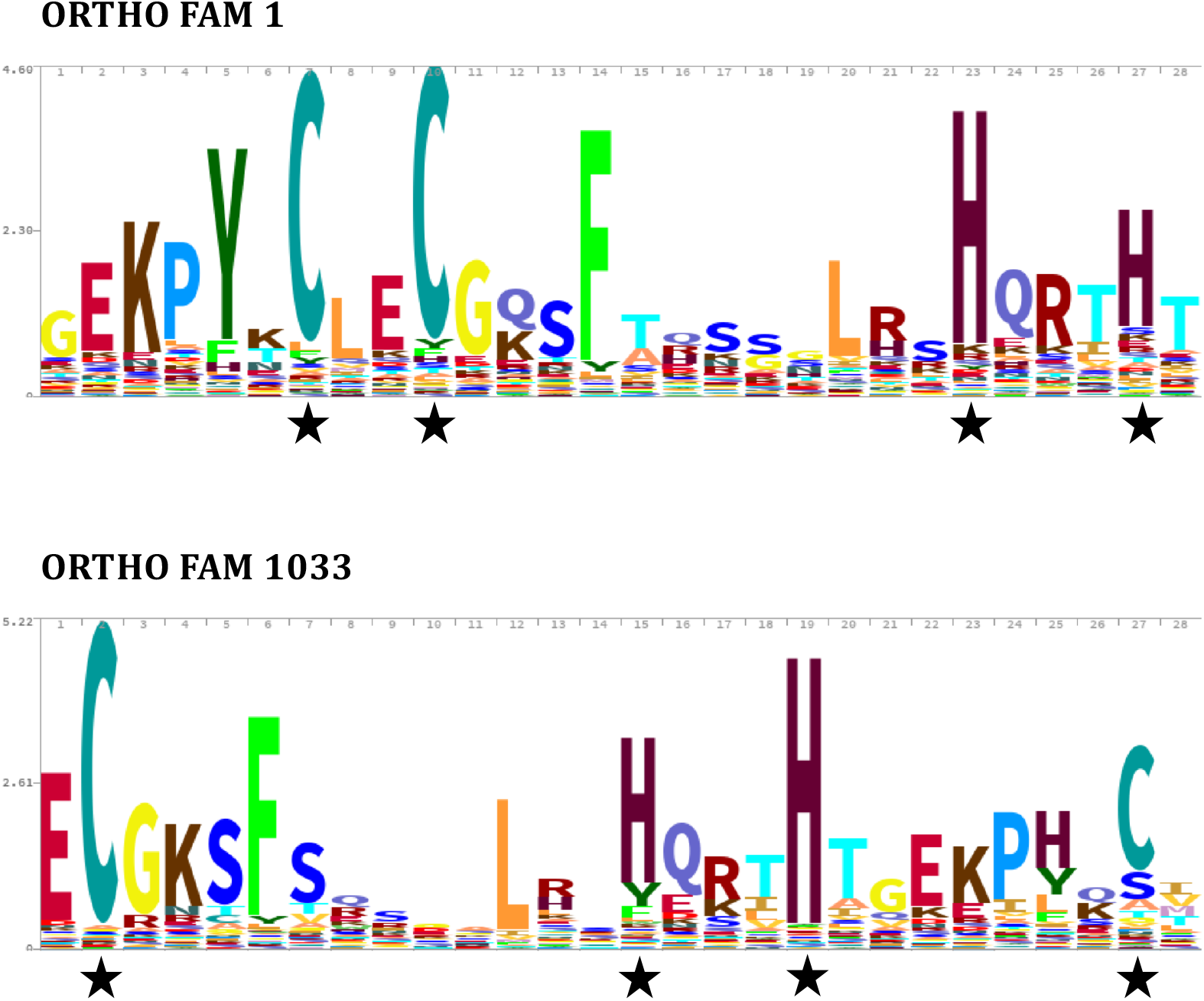
HMM of the two main zinc-finger protein families. These two families were phased differently and not merged. The two cysteine and the two histidine residues that are the hallmark of all zinc-finger proteins are indicated by black stars.

