## Supplementary material for "Megasatellite formation and evolution in vertebrates": Table 1

**Table 1: List and properties of the 62 genomes used in the present study**

| Species name | Common name | Clade | Genome size (1) | Gene number | ORTHO FAM number | Release date |
|---|---|---|---|---|---|---|
| *Ailuropoda melanoleuca* | Giant panda | Eutherian | 6,0E+06 | 19343 | 92 | 04-sept-18 |
| *Anas platyrhynchos* | Mallard (dabbling duck) | Aves | 6,0E+06 | 15634 | 29 | 04-sept-18 |
| *Anolis carolinensis* | Green anole | Lepidosauria | 1,1E+09 | 18595 | 45 | 05-sept-18 |
| *Astyanax mexicanus* | Mexican tetra (blind cave fish) | Teleostei | 9,4E+08 | 26698 | 41 | 05-sept-18 |
| *Bos taurus* | Cow | Eutherian | 2,7E+09 | 19994 | 79 | 05-sept-18 |
| *Callithrix jacchus* | Common marmoset | Primate | 2,1E+08 | 19690 | 87 | 05-sept-18 |
| *Canis familiaris* | Dog | Eutherian | 2,3E+09 | 19856 | 79 | 04-sept-18 |
| *Cavia porcellus* | Guinea pig | Eutherian | 8,9E+07 | 18095 | 70 | 04-sept-18 |
| *Chlorocebus sabaeus* | Green monkey | Primate | 2,8E+09 | 19165 | 106 | 05-sept-18 |
| *Choloepus hoffmanni* | Hoffmann's two-toed sloth | Primate | 6,3E+05 | 12393 | 53 | 05-sept-18 |
| *Danio rerio (2)* | Zebrafish | Teleostei | 1,3E+09 | 30313 | ND | 05-sept-18 |
| *Dasypus novemcinctus* | Nine-banded armadillo | Eutherian | 1,4E+07 | 22711 | 91 | 05-sept-18 |
| *Dipodomys ordii* | Ord's kangaroo rat | Eutherian | 6,8E+07 | 16911 | 62 | 05-sept-18 |
| *Echinops telfairi* | Lesser hedgehog tenrec | Eutherian | 1,8E+06 | 16575 | 67 | 04-sept-18 |
| *Equus caballus* | Horse | Eutherian | 2,4E+09 | 20449 | 72 (3) | 05-sept-18 |
| *Erinaceus europaeus* | European hedgehog | Eutherian | 1,6E+06 | 14601 | 57 | 04-sept-18 |
| *Felis catus* | Cat | Eutherian | 2,5E+09 | 19446 | 71 | 04-sept-18 |
| *Ficedula albicollis* | Collared flycatcher | Aves | 2,7E+07 | 15303 | 42 (3) | 04-sept-18 |
| *Gadus morhua* | Cod | Teleostei | 3,9E+06 | 20095 | 34 | 05-sept-18 |
| *Gallus gallus* | Chicken | Aves | 1,0E+09 | 18346 | 42 | 05-sept-18 |
| *Gasterosteus aculeatus* | Stickleback | Teleostei | 4,1E+08 | 20787 | 39 | 05-sept-18 |
| *Gorilla gorilla* | Gorilla | Primate | 2,9E+09 | 21794 | 109 | 04-sept-18 |
| *Homo sapiens* | Man | Primate | 3,1E+09 | 23358 | 121 | 05-sept-18 |
| *Ictidomys tridecemlineatus* | Squirrel | Eutherian | 5,8E+07 | 18474 | 68 | 05-sept-18 |
| *Latimeria chalumnae* | Coelacanth | Actinistia | 1,1E+07 | 19569 | 49 | 05-sept-18 |
| *Lepisosteus oculatus* | Spotted gar | Ginglymodi | 8,9E+08 | 18341 | 51 | 04-sept-18 |
| *Loxodonta africana* | Elephant | Eutherian | 1,3E+08 | 20033 | 86 | 04-sept-18 |
| *Macaca mulatta* | Macaque | Primate | 2,8E+09 | 21099 | 100 | 05-sept-18 |
| *Macropus eugenii (2)* | Wallaby | Marsupiala | 5,4E+05 | 15290 | ND | 23-nov-16 |
| *Meleagris gallopavo* | Turkey | Aves | 1,0E+09 | 14123 | 40 (3) | 05-sept-18 |
| *Microcebus murinus* | Mouse lemur | Eutherian | 2,4E+09 | 18895 | 87 | 05-sept-18 |
| *Monodelphis domestica* | Opossum | Marsupiala | 3,6E+09 | 21327 | 55 | 04-sept-18 |
| *Mus musculus* | Mouse | Eutherian | 2,7E+09 | 22969 | 133 (3) | 05-sept-18 |
| *Mustela putorius furo* | Ferret | Eutherian | 5,2E+07 | 19910 | 85 | 05-sept-18 |
| *Myotis lucifugus* | Microbat (little brown bat) | Eutherian | 6,5E+07 | 19728 | 80 | 05-sept-18 |
| *Nomascus leucogenys* | Gibbon | Primate | 2,8E+09 | 20794 | 99 | 05-sept-18 |
| *Ochotona princeps* | Pika | Eutherian | 2,6E+06 | 16006 | 76 | 04-sept-18 |
| *Oreochromis niloticus* | Tilapia | Teleostei | 1,4E+07 | 21437 | 40 | 04-sept-18 |
| *Ornithorhynchus anatinus* | Platypus | Monotreme | 4,6E+08 | 21698 | 32 | 04-sept-18 |
| *Oryctolagus cuniculus* | Rabbit | Eutherian | 2,3E+09 | 19293 | 76 | 05-sept-18 |
| *Oryzias latipes* | Medaka | Teleostei | 7,3E+08 | 23622 | 52 (3) | 05-sept-18 |

| | | | | | | |
|---|---|---|---|---|---|---|
| *Otolemur garnettii* | Bushbaby | Primate | 7,3E+07 | 19506 | 85 | 04-sept-18 |
| *Ovis aries* | Sheep | Eutherian | 2,6E+09 | 20921 | 96 | 05-sept-18 |
| *Pan troglodytes* | Chimpanzee | Primate | 3,0E+09 | 23534 | 110 | 04-sept-18 |
| *Papio anubis* | Olive baboon | Primate | 2,7E+09 | 21647 | 112 | 05-sept-18 |
| *Pelodiscus sinensis* | Chinese softshell turtle | Sauropsida | 1,6E+07 | 18189 | 39 | 05-sept-18 |
| *Petromyzon marinus* | Sea lamprey | Agnatha | 4,7E+06 | 10415 | 19 | 04-sept-18 |
| *Poecilia formosa* | Amazon molly | Teleostei | 7,4E+06 | 23615 | 50 (3) | 05-sept-18 |
| *Pongo abelii* | Orangutan | Primate | 3,1E+09 | 20424 | 98 | 05-sept-18 |
| *Procavia capensis* | Rock hyrax | Eutherian | 1,3E+06 | 16057 | 75 | 04-sept-18 |
| *Pteropus vampyrus* | Large fruit bat (megabat) | Eutherian | 1,9E+06 | 16990 | 84 | 05-sept-18 |
| *Rattus norvegicus* | Rat | Eutherian | 2,8E+09 | 22250 | 86 | 05-sept-18 |
| *Saccharomyces cerevisiae* | Baker's yeast | Fungi | 1,2E+07 | 6600 | 1 | 05-sept-18 |
| *Sarcophilus harrisii* | Tasmanian devil | Marsupiala | 5,3E+06 | 18788 | 45 | 05-sept-18 |
| *Sorex araneus* | Shrew | Eutherian | 2,3E+06 | 13187 | 48 | 05-sept-18 |
| *Sus scrofa* | Pig (wild boar) | Eutherian | 2,4E+09 | 22452 | 89 | 04-sept-18 |
| *Taeniopygia guttata* | Zebra finch | Aves | 1,2E+09 | 17488 | 35 | 04-sept-18 |
| *Tarsius syrichta (2)* | Tarsier | Primate | 1,2E+06 | 13628 | ND | 24-nov-16 |
| *Tetraodon nigroviridis* | Pufferfish | Teleostei | 3,5E+08 | 19602 | 38 | 05-sept-18 |
| *Tursiops truncatus* | Dolphin | Eutherian | 2,2E+06 | 16550 | 87 | 05-sept-18 |
| *Vicugna pacos* | Alpaca | Eutherian | 5,5E+06 | 11765 | 49 | 04-sept-18 |
| *Xenopus tropicalis* | Xenopus (western clawed frog) | Amphibia | 7,8E+06 | 18442 | 39 | 05-sept-18 |

(1) Genome size was calculated by adding all contig lengths.
(2) These three species were originally included in the first analysis on total genomic DNA but were dropped in the second analysis on protein sequences (see text for details).
(3) Megasatellites are more frequent than expected number (p-value < 0.01) (see text for details).
