## Supplementary material for "Megasatellite formation and evolution in vertebrates": Table S1

**Supplemental Table S1: Number and percentage of each megasatellite family**

| Megasatellite type | Number (%) |
|---|---|
| **Monomega** | 65 (45.8%) |
| **Multimega** | 18 (12.7%) |
| **Hidden multimega** | 28 (19.7%) |
| **Overmega** | 26 (18.3%) |
| **Mixmega** | 4 (2.8%) |
| **Intromega** | 1 (0.7%) |
| **Total** | **142** |
