## Supplementary material for "Megasatellite formation and evolution in vertebrates": Table S2

**Supplemental Table S2: Number of families represented in each clade**

| Clade | Number |
|---|---|
| Agnatha | 17 |
| Ginglymodi | 47 |
| Teleostei | 55 |
| Actinistia | 44 |
| Amphibia | 37 |
| Sauropsida | 35 |
| Lepidosauria | 43 |
| Aves | 45 |
| Monotreme | 30 |
| Marsupiala | 60 |
| Eutherian | 107 |
| Primate | 116 |
| Fungi | 1 |
