## Supplementary material for "Megasatellite formation and evolution in vertebrates": Table S3

**Supplemental table S3: List and repeat unit number of polyubiquitin genes**

| Clade | Species | Nbr of repeats | Nbr of repeats in [1] |
|---|---|---|---|
| **Actinistia** | *Latimeria chalumnae* | 2 | |
| **Agnatha** | *Petromyzon marinus* | 5 | |
| | *Petromyzon marinus* | 6 | |
| **Teleostei** | *Gadus morhua* | 2 | |
| | *Poecilia formosa* | 2 | |
| | *Poecilia formosa* | 3 | |
| | *Poecilia formosa* | 4 | |
| | *Tetraodon nigroviridis* | 3 | |
| | *Astyanax mexicanus* | 10 | |
| | *Gasterosteus aculeatus* | 18 | |
| | *Oryzias latipes* | 12 | |
| **Lepidosauria** | *Anolis carolinensis* | 6 | |
| **Sauropsida** | *Pelodiscus sinensis* | 7 | |
| **Aves** | *Taeniopygia guttata* | 3 | |
| | *Gallus gallus* | 4 | 4 |
| | *Gallus gallus* | 5 | |
| | *Ficedula albicollis* | 5 | |
| | *Ficedula albicollis* | 5 | |
| **Marsupiala** | *Monodelphis domestica* | 4 | |
| | *Sarcophilus harrisii* | 4 | |
| | *Sarcophilus harrisii* | 5 | |
| **Eutherian** | *Ailuropoda melanoleuca* | 4 | |
| | *Bos taurus* | 5 | |
| | *Bos taurus* | 5 | |
| | *Canis familiaris* | 3 | |
| | *Cavia porcellus* | 3 | |
| | *Cavia porcellus* | 4 | |
| | *Dasypus novemcinctus* | 3 | |
| | *Dasypus novemcinctus* | 6 | |
| | *Dipodomys ordii* | 3 | |
| | *Equus caballus* | 4 | |
| | *Felis catus* | 4 | |
| | *Loxondata africana* | 5 | |
| | *Loxondata africana* | 6 | |
| | *Microcebus murinus* | 4 | |
| | *Mus musculus* | 5 | 4 |
| | *Mus musculus* | 10 | 9 |
| | *Mustela putorius furo* | 5 | |
| | *Myotis lucifugus* | 2 | |
| | *Myotis lucifugus* | 5 | |
| | *Ochotona princeps* | 2 | |
| | *Ochotona princeps* | 4 | |
| | *Oryctolagus cuniculus* | 3 | |
| | *Ovis aries* | 4 | |

| | | | |
|---|---|---|---|
| | *Ovis aries* | 16 | |
| | *Procavia capensis* | 9 | |
| | *Pteropus Vampyrus* | 4 | |
| | *Rattus norvegicus* | 2 | |
| | *Rattus norvegicus* | 11 | |
| | *Sus scrofa* | 4 | |
| | *Sus scrofa* | 12 | |
| | *Tursiops truncatus* | 4 | |
| | *Tursiops truncatus* | 7 | |
| **Primate** | *Callithrix jacchus* | 3 | |
| | *Chlorocebus sabaeus* | 3 | |
| | *Chlorocebus sabaeus* | 4 | |
| | *Gorilla gorilla* | 3 | |
| | *Gorilla gorilla* | 4 | |
| | *Gorilla gorilla* | 5 | |
| | *Homo sapiens* | 8 | 3 |
| | *Homo sapiens* | 20 | 11 |
| | *Macaca mulatta* | 4 | |
| | *Macaca mulatta* | 4 | |
| | *Nomascus leucogenys* | 3 | |
| | *Nomascus leucogenys* | 5 | |
| | *Otolemur garnettii* | 3 | |
| | *Pan troglodytes* | 3 | |
| | *Pan troglodytes* | 4 | |
| | *Pan troglodytes* | 10 | |
| | *Papio anubis* | 3 | |
| | *Papio anubis* | 3 | |
| | *Papio anubis* | 5 | |
| | *Pongo abelii* | 4 | |
| | *Pongo abelii* | 9 | |
| **Fungi** | *Saccharomyces cerevisiae* | 6 | 6 |

1. Gemayel, R. *et al.* Variable repeats in the eukaryotic polyubiquitin gene ubi4 modulate proteostasis and stress survival. *Nat. Commun.* **8**, 397 (2017).
